# Eco-evolutionary dynamics are shaped by competition in experimental range expansions

**DOI:** 10.64898/2026.06.01.729372

**Authors:** Carla A. Urquhart, Takuji Usui, Amy L. Angert, Jennifer L. Williams

## Abstract

Most theory and empirical research on range expansion assumes populations spread into empty landscapes with abundant resources, however expanding populations are likely to compete with residents. In mathematical models, interspecific competition can lead to pushed wave dynamics, where expansions are driven mainly by individuals dispersing from the core, leading to steeper wavefronts and increased genetic diversity at the edge. These predictions are yet to be tested empirically, and the role of interspecific competition in mediating evolution during range expansion is unclear. We used an experimental system with two duckweed species to ask if interspecific competition leads to pushed-like dynamics and to assess how competition alters evolution during range expansion. We found that competition with a resident reduced expansion speed and absolute variance among replicate expansions, suggesting competition makes expansion speed more predictable. Interspecific competition also changed the relative frequencies of genotypes at the leading edge. While competition was associated with some features of pushed waves, genotype diversity did not vary between treatments. Our results demonstrate that demographic and evolutionary patterns associated with pushed waves may not be universal, and that incorporating selective pressures into future research on eco-evolutionary dynamics of range expansion is key to understanding spreading populations in nature.

## Introduction

With increasing environmental change, many species ranges are shifting and expanding (Nadeau and Urban 2019), making understanding the processes underpinning ecological and evolutionary dynamics of range expansion key to predicting contemporary population dynamics. In a large majority of both theory (Clark et al. 2001; Phillips 2009, 2015; Burton et al. 2010) and experiments (Fronhofer and Altermatt 2015; Williams et al. 2016a; Ochocki and Miller 2017; Weiss-Lehman et al. 2017; Ochocki et al. 2019; Weiss-Lehman et al. 2019; Deshpande and Fronhofer 2022; Usui and Angert 2024) it is assumed that populations spread into empty landscapes with abundant resources. However, these conditions are unlikely to be experienced by expanding species in nature (Gilman et al. 2010). Interspecific interactions, and competition in particular, may fundamentally alter the ecological and evolutionary processes occurring at the leading edge of expanding populations.

In general, selection at the leading edge is predicted to favour traits related to reproduction and dispersal, resulting in a process often termed spatial selection (Phillips 2009; Burton et al. 2010; Van Petegem et al. 2016; Phillips and Perkins 2019). While spatial evolutionary processes are thought to be ubiquitous, their dominance relies on the assumptions that: 1) leading-edge populations are small, and 2) the environment at the leading edge is more suitable than at the core due to an abundance of resources with fewer individuals to utilise them. When these conditions are not met, spatial evolutionary processes such as spatial selection and founder effects are less likely to occur. This has been shown in a number of systems with moderate to strong selective pressures, where more adaptive traits are favoured, such as with environmental gradients (Hargreaves and Eckert 2014; Garnier and Lewis 2016; Usui and Angert 2024) and landscape fragmentation (Williams et al. 2016a, 2016b; Urquhart and Williams 2021) as well as with high mutation rates (Poloni and Lutscher 2023). Interspecific competition can result in slower expansion rates (Okubo et al. 1989; Legault et al. 2020; McHugh et al. 2025) and instead may create stronger selection pressure for traits related to competition.

Differences in leading edge evolutionary processes due to competition may also alter variability in expansion speed among replicates in experimental expansions, a useful proxy for predictability in nature (Phillips 2015). In experimental range expansions with evolving and non-evolving populations, evolution has been found to modify variability in speed but not in a consistent way (Williams et al. 2019). In experiments (Usui and Angert 2024) and simulations (Benning et al. 2024) with environmental gradients, where expansions are slowed due to strong selective processes, variability between replicates was higher than in more benign environments. These results contradict predictions that adaptive evolutionary processes would make experimental populations more genetically similar, and thus expansions less variable. It is important to note, however, that all of these studies use metrics based on the coefficient of variation to quantify variability, which controls for differences in mean speed. This allows us to assess differences in relative variability, and to better understand underlying processes. However, measures of absolute variance may be more appropriate for use as a proxy for predictability as we are interested in absolute error, or the difference between predicted and realized expansion speed.

Competitive interactions can also result in what are termed “pushed” expansions (Lewis et al. 2002), which can occur when fitness declines at the leading edge of an expanding population (Miller et al. 2020). In mathematical models, pushed waves are a result of positive density dependence in population growth or dispersal (Bonnefon et al. 2013, 2014; Birzu et al. 2019), or environmental pressures at the leading edge, such as temperature gradients (Garnier and Lewis 2016) or landscape fragmentation (Wang et al. 2019). A common biological interpretation is that pushed expansions are driven by individuals dispersing from a higher fitness core population, as opposed to being “pulled” forward by dispersers from the leading edge (Bonnefon et al. 2013; Miller et al. 2020). This can alter the shape of the expanding wave, as pulled wave expansions extend more quickly over space at low densities than pushed wave expansions, which are characterized by a steeper wave front (Figure 1 b,c) and increased population sizes at the leading edge (Williams et al. 2019; Miller et al. 2020). But, it is important to note that wave form dynamics (i.e. whether an expansion is pushed or pulled) are fundamentally a property of mathematical models, and difficult to map to biology. That is, pushed and pulled waves arise due to mathematical properties of the dynamical equations used to model them, and biological interpretations for these properties remain somewhat elusive. In highly stochastic systems it can be difficult to determine if an expansion is really “pulled” or “pushed.” However, we can look for properties that distinguish each type (such as wave steepness and leading edge population densities), and for positive density dependence in reproduction or dispersal that would reduce the selective advantage of dispersal and low density fecundity at the leading edge.

**Fig 1.**
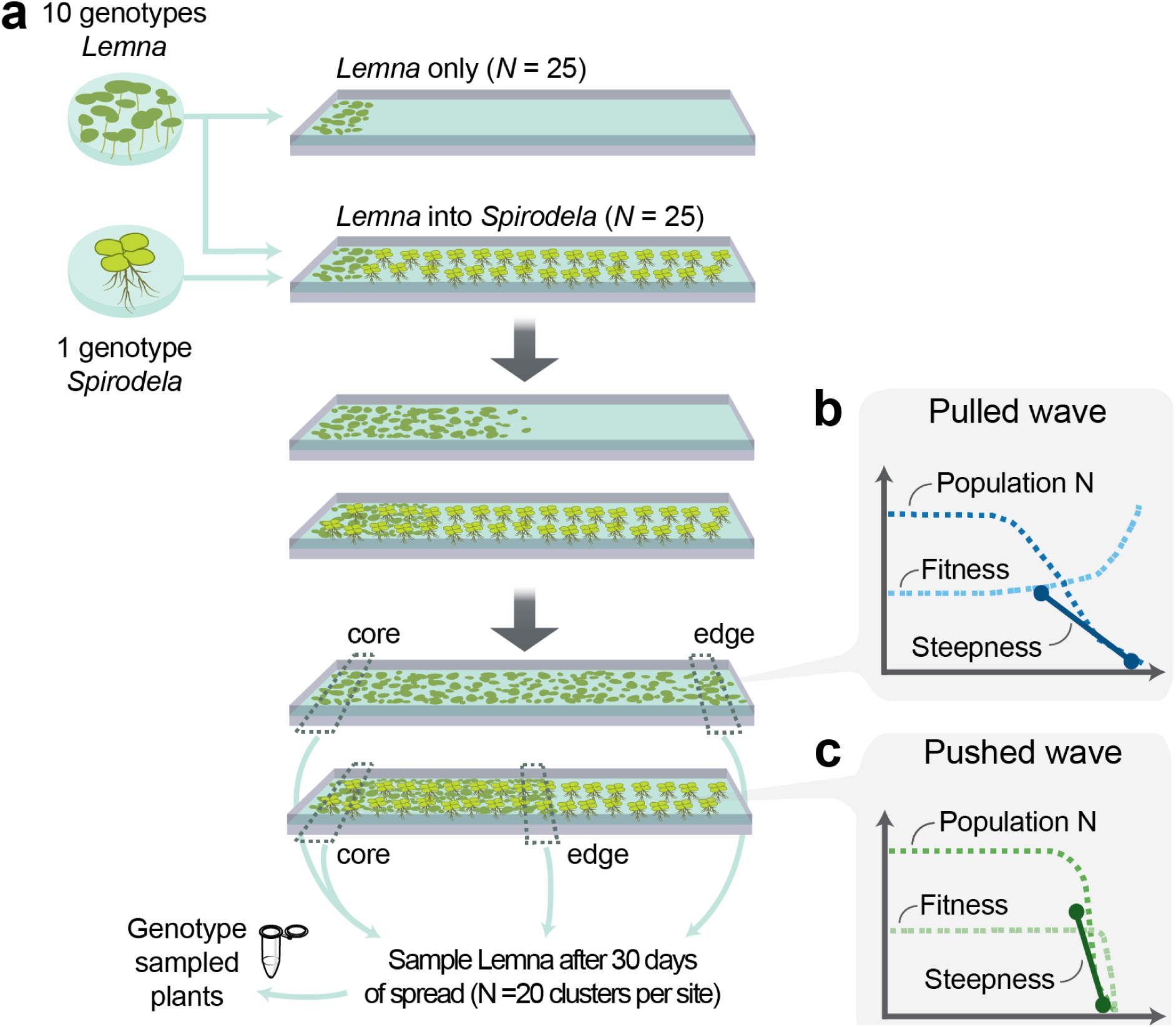
Experimental design and predictions for experimental range expansions in linear mesocosms with two species of duckweed. (a) Illustration of one experimental mesocosm per treatment. (b,c) Predictions for wave steepness for pulled (b) and pushed (c) waves. We expected the expansions with *Spirodela* would spread as pushed waves due to lower fitness of leading edge individuals. We measured wave steepness, the distance between the leading edge and where population densities reached 50% of peak density, as an estimate of whether expansions were more pushed or pulled.

Features of pushed waves, such as higher leading-edge population density, mean they may exhibit very different evolutionary dynamics from pulled wave expansions that have traditionally been considered in eco-evolutionary theory. In addition to spatial selection, spatial evolutionary processes that occur at the leading edge are expected to be largely dominated by stochastic processes resulting from serial founder events (Excoffier and Ray 2008; Peischl et al. 2015; Peter and Slatkin 2015). When stochastic processes are dominant, a population’s expansion speed may be more variable, and thus less predictable, as key traits such as reproduction and dispersal can vary widely between replicate populations (Williams et al. 2019). In pushed expansions, individuals from the range core are more likely to disperse to and accumulate at the leading edge, increasing gene flow and thus reducing the role of founder effects and facilitating adaptation at the leading edge (Miller et al. 2020, Burton et al. 2010). In mathematical models and experiments, pushed wave expansions have been shown to retain more genetic diversity at the leading edge than expected (Garnier and Lewis 2016; Dahirel et al. 2021), potentially making expansion speed more predictable. With weaker founder effects, spatial genetic structure across the expanding population may also be less likely to develop. That is, with competition from a resident species, leading edge populations may be more genetically similar to core populations than when competition is not present.

Thus, we can predict that competition may result in higher genetic diversity at the leading edge as in other instances of pushed wave dynamics. In natural populations, a limited role for stochastic evolutionary processes and spatial selection due to competition could mean more genetically diverse, locally adapted and resilient populations at longer time scales after range expansions or shifts. As we expect that most species are likely to be impacted by interspecific competition with resident communities, it is crucial to develop a better understanding of the spatial evolutionary processes that might occur under these conditions.

To examine the role of a resident competitor in shaping the dynamics of expansion and evolution in a range-expanding species, we conducted experimental expansions using a two-species duckweed system in a greenhouse mesocosm array. Duckweeds are an ideal model system for range expansions, as they are short-lived, with simple life histories and have an extremely low mutation rate, making evolution readily quantified by tracking genotype frequencies over time and space (Hart et al. 2019; Acosta et al. 2021; Usui and Angert 2024). Mesocosm experiments allow us to assess population dynamics and evolutionary trajectories while maintaining strong links to theory (Lustenhouwer et al. 2023). Here, we allowed experimental populations of common duckweeds in the *Lemna* species complex to spread into environments with ample resources and no competition as well as into environments with an established population of the greater duckweed, *Spirodela polyrhiza*. We asked, how does competition with a resident species alter expansion speed and the expanding wave form dynamics (i.e., pushed or pulled waves), as well as variability in speed and population densities? And, how does competition impact evolutionary processes at the leading edge and the development of spatial genetic structure across expanding populations? We hypothesized that expansions with a resident competitor would be slower, with less variability in speed among replicate populations than those spreading in empty space because expanding populations facing interspecific competition at the leading edge could have more time to grow to higher densities as the leading edge advances more slowly. This may in turn increase the role of natural selection for traits related to competitive ability, such as specific leaf area and root structures, and reduce variability in both speed and in traits found at the leading edge. As theory predicts that range expansions into competitive environments will advance as pushed waves, we predicted higher population densities and genetic diversity at the leading edge with competition than in expansions into empty habitat.

## Methods

### Study system

Duckweeds are a globally distributed freshwater plant species that has been used increasingly as a model system in ecological and evolutionary research (Laird & Barks 2018). Their small size, very low rate of outcrossing and fast generation times, averaging 2-5 days under optimal conditions (Landolt 1986), make this an ideal system for mesocosm experiments. Clonal growth occurs through the budding of individual, offspring fronds from the parental frond, with offspring fronds staying connected to the parental frond in clonal ramets (clusters) before their eventual release. We assembled populations using 10 unique genotypes of *Lemna* spp. sampled from southwestern British Columbia (BC), Canada, and northwestern Washington, USA (Usui and Angert 2024). Genetic barcoding conducted previously indicated that these genotypes come from two *Lemna* species, with 6 genotypes of *Lemna minor* and 4 genotypes of *Lemna japonica*, a hybrid of *Lemna minor* and *Lemna turionifera*. As these species are very similar genetically and ecologically, we consider them together as the *Lemna* species complex (Braglia et al. 2021; Usui and Angert 2024). We used the closely related, but morphologically and ecologically distinct *Spirodela polyrhiza* as the resident species, which is basal on the phylogenetic tree to all other duckweed species (Tippery and Les 2020)*. Spirodela polyrhiza* populations used in the experiment consisted of a single genotype collected from a pond in Vancouver, BC, in order to exclude effects of evolution in resident populations.

### Experimental range expansions

We conducted experimental range expansions in populations of *Lemna* with and without interspecific competition using gutter mesocosms in a greenhouse at the University of British Columbia (Fig. 1). Fifty experimental mesocosms were constructed from aluminum gutters (150 × 12.5 cm; L × W) and placed across five benches. Gutters were filled with a low nutrient media solution (Docauer 1983) and treated with black pond dye (Natural Waterscapes, Pennsylvania USA) to limit algal growth. Each mesocosm was randomly assigned to the competition (+*Spirodela*) or control (-*Spirodela*) treatment. Competitive mesocosms were seeded with populations of *S. polyrhiza* evenly distributed throughout the gutter, save the first 10 cm of each gutter, at an approximate density of 10 fronds per centimeter measured along the width of the gutter, 3 days before initializing the experiment. On day 1 of the experiment, 200 fronds of an equal frequency of 10 *Lemna* genotypes were placed along the first 8 cm of each gutter. To account for disturbances to water movement during frond placement, fronds that floated beyond the 8 cm core were relocated each day until day 3, unless the whole population floated to the opposite end of the gutter, in which case they were left to spread in the opposite direction. On day 4 the location of population cores was recorded, which we defined as the 8 cm occupied by *Lemna* nearest to the gutter end. Most cores were designated as the same 0-8 cm as populations were initiated in, however one population floated to the opposite side of the gutter and 5 populations settled outside the initial 0-8 cm range, but still near the gutter end. Two populations that did not reach the gutter end by day 7 were excluded from analyses, as they did not have a single leading edge. As *S. polyrhiza* at high densities acts as a physical barrier to spread of *Lemna*, *S. polyrhiza* populations were thinned on day 10 to maintain consistent densities across replicates and over time. Surface algae was removed from gutters as necessary to allow populations to disperse over the water surface. Populations spread for 30 days. On the final day, 20 distinct ramets were sampled from the leading edge (farthest 8 cm from the core) and core (as established for each replicate on day 4) of each replicate for genotyping. An average of 12 distinct samples per replicate per location were successfully genotyped, and eight replicates with fewer than five genotyped individuals at the edge and core were dropped from all analyses. In addition to the ten replicate mesocosms mentioned above that were dropped, two replicates were dropped due to an irrigation system failure during the experiment and five were dropped due to gutters being jostled during data collection. In all, we retained 33 replicate mesocosms, 14 without *Spirodela polyrhiza* and 19 with *Spirodela* as the resident competitor (hereafter, -*Spirodela* and +*Spirodela* respectively).

### Spatial population dynamics and expansion speed

Expansion speed and population densities were estimated along the length of experimental ranges using image analysis. We photographed the extent of each replicate expansion every 3-4 days, and recorded the position of the furthest forward frond. Expansion extent was used as a proxy for average expansion speed, as speed is highly variable across time in our experiment. Extent was measured as the distance in centimeters from the furthest forward frond and the population core, as established in the first week of the experiment. To estimate population density at the end of the experiment, we counted the number of fronds along linear transects in ImageJ (Schneider et al. 2012). We drew cross-sectional transects perpendicular to the direction of expansion starting from the middle of the identified core, and then every 10 cm until 15 cm behind the furthest forward frond, where we increased the transect density to every 2 cm. All *Lemna* fronds, excluding dead ones, intersecting the transects were counted. We then calculated a metric for wave steepness using a method based on Williams et al. (2019) as a way to estimate whether expansions spread as pushed or pulled waves (Fig 1). Wave steepness (*S*) is defined as the distance between the furthest forward frond and the transect closest to the leading edge with a population density greater than 50% of the largest population density observed in the same replicate:

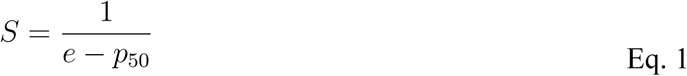

where *e* is the location of the furthest forward individual, *p_50_* is the furthest forward location to reach 50% of the maximum population density. When *p_50_* was equal to the edge distance, wave steepness was set to 1 as a maximum steepness to avoid undefined values. While this does not define pushed and pulled waves using the mathematical criteria established in analytical models (Lewis 2000; Bonnefon et al. 2013), it provides a useful proxy as most biological features of pushed waves are thought to be driven by relatively higher leading edge population densities (Miller et al. 2020). To compare among-replicate variability in expansion extent and wave steepness between experimental treatments, we used two metrics: the log variation ratio test (lnVR) and the log coefficient of variation ratio test (lnCVR) (Nakagawa et al. 2015; Usui and Angert 2024) to assess absolute and relative variability, respectively:

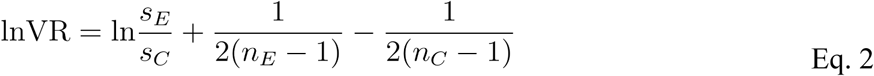

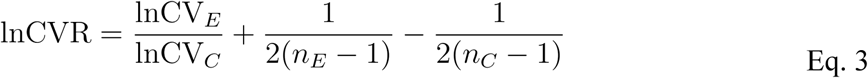

which are the ratio of the standard deviations (Eq. 2) and coefficients of variation (Eq. 3) in the experimental (*E*) and control (*C*) treatments – in this case, + and -*Spirodela –* with correction terms for sample sizes *n_E_* and *n_C_*. In both metrics, variability is higher in the experimental (+*Spirodela*) treatment when the result is positive, and lower if negative. The significance of the results of both metrics was assessed using 95% bootstrapped confidence intervals, where results are not significant when the interval includes zero. We analyzed differences in extent and wave steepness between treatments using analysis of variance in R 4.5.1 (R Core Team 2025).

### Evolutionary dynamics

To assess differences in genetic diversity and evolutionary trajectories at the leading edge, we compared relative frequencies of the ten genetically distinct lineages of *Lemna* species used in the experiment (referred to as genotypes hereafter). We defined evolution in our experiment as changes in either genotype frequencies or genotype-weighted trait values from standing variation. Mutation is unlikely to have contributed to evolutionary change in our experiment due to the short experimental duration and low mutation rate of *Lemna* species (Sandler et al. 2020). To identify genotypes, DNA was extracted from samples collected from the edge and core of experimental mesocosms on day 30 using a CTAB extraction protocol (Healey et al. 2014), and samples were genotyped using microsatellite analysis with 4 markers in a single multiplex reaction (Hart et al. 2019, see <u>Appendix S1</u>). We retained all replicates that had at least 5 successfully genotyped individuals from the edge and core, although some replicates had up to 20 genotyped individuals. For analyses of genetic diversity, we used a subset of 27 replicate expansions (n=13 -*Spirodela* and n=14 +*Spirodela)* that had at least 10 genotyped individuals from both the edge and core.

In addition to genotype frequencies at the leading edge and core, we assessed differences in genetic diversity and genotype-weighted trait means for selected traits expected to be related to competitive ability. Genotype diversity in edge and core samples was calculated using inverse Simpson’s diversity index where “species” are genotypes, using the vegan package in R (Okansen et al. 2025). We chose Simpson’s index as we were interested in whether a single or few species were dominant, and this index should be less sensitive to the upper bound on “richness” imposed by our experiment than other simple diversity metrics (DeJong 1975). Simpson’s diversity is also analogous to heterozygosity, a metric commonly used in population genetics, but is calculated at the level of the individual, rather than the allele. We assessed differences in diversity between treatments and between edge and core populations using analysis of variance in R (R v. 4.5.1, R Core Team 2025). To compare diversity in our samples to the starting populations, we constructed bootstrapped confidence intervals on mean diversity. Where these intervals overlapped 10 there was no difference in diversity from initial populations, as the inverse Simpson’s index for a population with equal frequencies of 10 types is also 10. Changes in competition-related traits were assessed by calculating genotype-weighted trait means for four traits shown in the literature to be related to competitive ability in duckweeds (Hart et al. 2019). We measured root length and reproduction (per capita growth) in 9 fronds from each genotype in laboratory microcosms under conditions as close as possible to those in the experiment, and calculated normalized trait means using z-scores to minimize the effect of different measurement units and magnitude of trait values across traits. We supplemented this with trait data previously collected for our experimental genotypes (Usui and Angert 2024) for specific leaf area and ramet size, which were also normalized across traits (see <u>Appendix S2</u>). We used linear models in R to test the effect of these measured traits on expansion extent (R v. 4.5.1, R Core Team 2025).

We compared the differences in genotype frequencies at the leading edge to both the initial frequencies (equal frequencies of 0.1) and between the edge and core at the end of the experiment. To assess the significance of our results, we constructed probability distributions for neutral genotype and trait variation to assess the probability of obtaining our results by chance. We simulated sampling from the initial genotype frequencies (1000 simulated samples for each sample size we obtained) and constructed 95% confidence intervals for genotype and trait frequencies from initial populations, applying a Bonferroni correction to account for comparisons across replicates as well as genotypes / traits. We also took pair-wise differences between 1000 pairs of simulated samples to construct 95% confidence intervals for differences in genotype frequencies and trait values between leading edge and core samples. When replicates fall outside of these intervals, it indicates that the values observed are unlikely to be due to random sampling from populations with the initial genotype frequencies. This method allowed us to assess the significance of observed evolutionary changes with variable sample sizes across replicates. In addition, we constructed bootstrapped 95% confidence intervals on the mean differences between edge and core for each genotype and trait.

## Results

### Effects of competition on expansion speed, variability and wave form dynamics

In line with our predictions, expansions in mesocosms with *Spirodela* were slower over the course of the experiment (Fig. 2), with significant differences in expansion extent observed consistently beginning on day 6 of the experiment, that is, after 1-2 generations of spread (F_1, 31_=25.24, p<0.001). These increased in magnitude until the end of the experiment on day 30 (F_1, 31_=78.61, p<0.001). We observed that populations in *-Spirodela* mesocosms were more likely to have pulsed expansions where a few fronds dispersed long distances beyond the leading edge, but then died or floated backward before reproducing (38% of expansions without competition, 0 with a resident competitor).

**Fig. 2.**
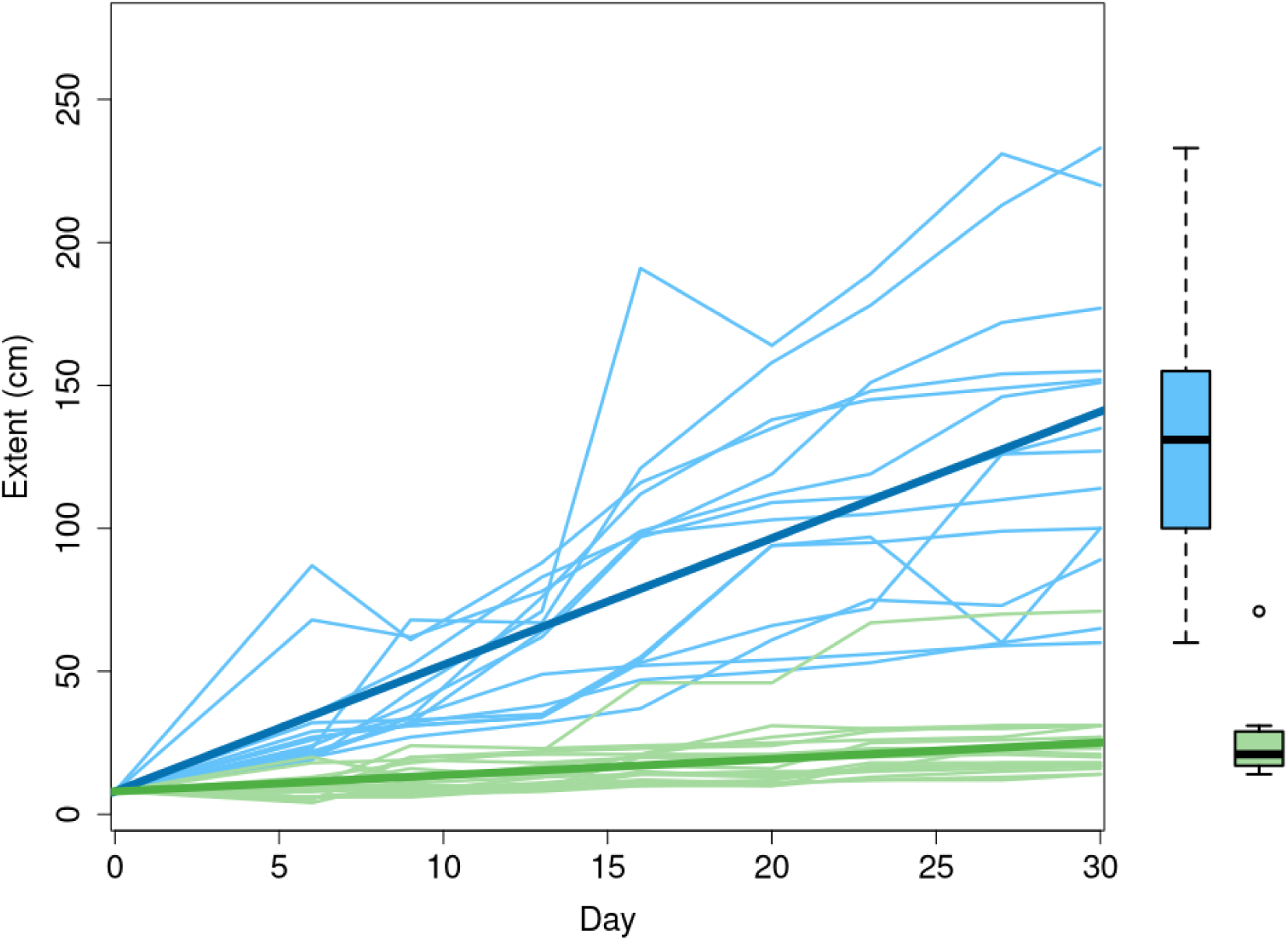
Expansion extent over time. Thin lines are individual replicates, thick lines are the modelled mean extents. Mesocosms with and without *Spirodela polyrhiza* are shown in green and blue, respectively. Boxplots on the right show the distribution of extents at the end of the experiment (Day 30).

While absolute variance and standard deviations of expansion extent were higher in mesocosms without *Spirodela* (lnVR = -1.41, 95% CI [-2.285, -0.501]), *+Spirodela* mesocosms had higher relative variation after controlling for differences in mean expansion extent, although this was not significant as assessed by overlap of the 95% CI with zero (lnCVR = 0.28, 95% CI [-0.4583, 1.0820]). High CV of extent in *+Spirodela* mesocosms was driven by one extreme outlier in the competitive treatment. When excluded, mesocosms without *Spirodela* have more variability in speed using both metrics but the difference remained non-significant using lnCVR (lnVR = -2.12, 95% CI[-2.546, -1.696]; lnCVR = -0.32, 95% CI [-0.7215, 0.0791]). This outlier replicate did not differ significantly from other replicates in its wavefront steepness, frequencies of genotypes or traits, nor was anything unusual observed in that replicate to justify its exclusion (see <u>Appendix S3</u>), therefore it was retained in the dataset.

Mesocosms with *Spirodela* had steeper wavefronts (F_1, 31_ = 29.06, p<0.001), measured as the distance between the leading edge and the furthest forward transect with at least 50% of the peak population density, with more absolute variability (lnVR = 0.62, 95% CI [-2.185, -0.428]). Relative variability in steepness was lower with *Spirodela,* but this difference was not significant (lnCVR = -0.33, 95% CI [-0.3664, 0.9892]). Contrary to our predictions, leading edge population densities did not differ between treatments (F_1,31_ = 0.189, p=0.66), and instead the observed differences in wave steepness were mostly driven by lower core population densities in *+Spirodela* mesocosms (F_1,31_ = 21.19, p < 0.001, and see Fig. 3).

**Fig. 3.**
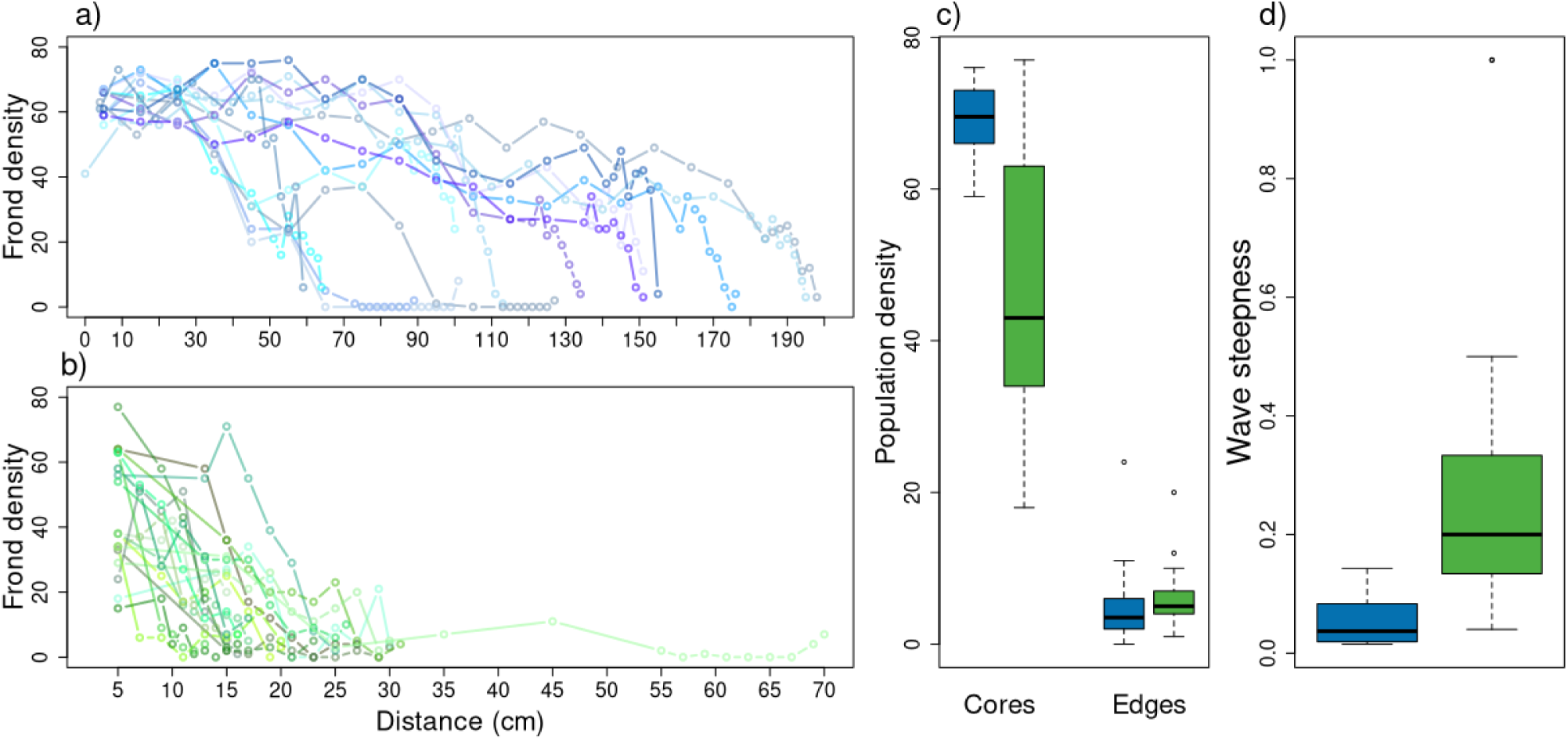
Population densities and wave steepness on day 30. (a,b) Density of fronds along cross-sections of each mesocosm are shown for mesocosms *-Spirodela* (a) and *+Spirodela* (b). Lines are individual replicates, points are cross-sections where densities were recorded. (c,d) Boxplots show the distribution of core and edge densities in each treatment (c) and wave steepness (d), a metric calculated using the distance between the furthest forward frond and the first cross-section to reach 50% of the maximum observed population density. Mesocosms with and without *Spirodela* are shown in green and blue, respectively.

### Effects of competition on genetic differentiation and genetic diversity

We found that LJ03 was the most common genotype in population cores and leading edges with *Spirodela*, while without *Spirodela* LJ04 was most common at the leading edge when genotype frequencies were pooled across replicates (Fig 4A-B). There was considerable variation in leading edge genotype frequencies between replicates overall, resulting in mostly non-significant differences in frequency between treatments (Fig. 4C). However, many of the non-significant replicates are genotype frequencies of zero where lineages went extinct. Where genotype frequencies increased significantly from initial conditions based on simulated sampling distributions (see Methods), genotype LJ04 was found at higher frequency at the leading edge without *Spirodela* than other genotypes (n = 4 of 9 total replicates with significant increases in genotype frequency), while in the + *Spirodela* mesocosms, no one genotype increased significantly more often than any others. Generally, we found that genotype-weighted traits at the leading edge did not vary between treatments (Fig. 4D). We found a significant positive correlation between one leading edge trait, root length, and expansion extent (β = 61.433 , t = 2.782 p = .0094) that was stronger in - *Spirodela* mesocosms as shown by a marginally significant interaction with treatment (F_1,31_ = 3.585, p = .0683), but there were no other significant correlations between traits and expansion speed (Fig. 6, and see Table S3).

**Fig 4.**
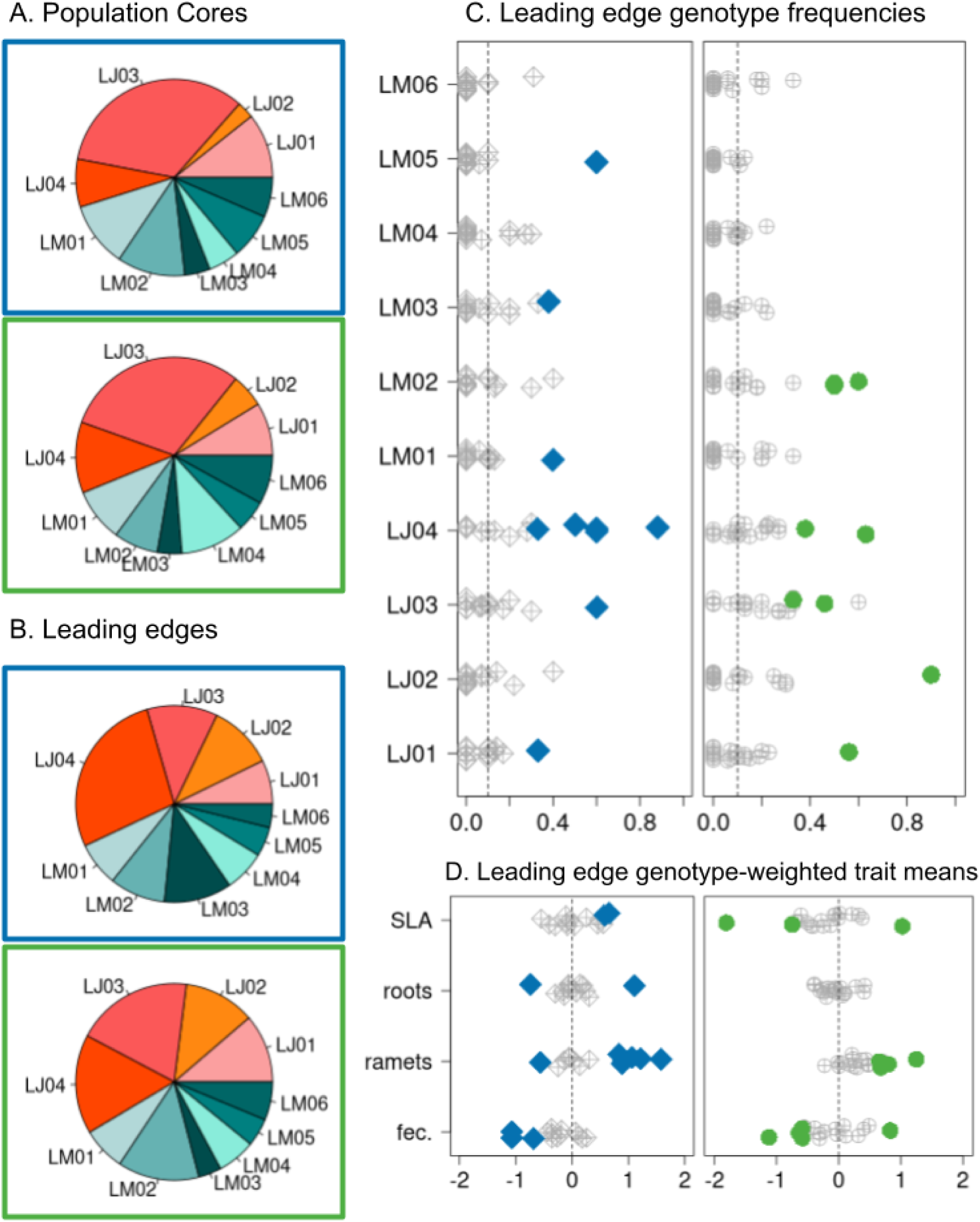
Leading edge genotype frequencies and traits. (a-b) Frequencies of each genotype on day 30, pooled across replicates for population cores (a) and leading edges (b). Charts framed in blue are *-Spirodela* mesocosms, and those framed in green are *+Spirodela*. Genotypes of *Lemna japonica* are shown in shades of orange, and *Lemna minor* are shades of teal. (c) Genotype frequencies and (d) genotype weighted trait means are shown by replicate at the leading edge on day 30. Points correspond to individual replicates. When values fall outside of simulated 95% confidence intervals for random sampling, *-Spirodela* mesocosms are shown in blue and *-Spirodela* in green, indicating significant change in frequency of that genotype or trait. Points are in grey otherwise. Dashed vertical lines show initial frequencies. Traits shown in (d) are, from top to bottom, specific leaf area (SLA), root length (roots), number of fronds per ramet (cluster) and low density population growth rate (fec.).

Genotype frequencies were rarely significantly different between the edge and core (freq. significant= 0.109, or n = 36 significant / 330 genotype x replicate combinations), but were significantly different more often in replicates without *Spirodela* (-*Spirodela* freq. significant = 0.15, +*Spirodela* freq. significant = 0.079, and see Fig 5A). In *-Spirodela* mesocosms, two genotypes were significantly more common at the leading edge than at the core (LJ04: mean freq. difference = 0.20, 95% CI [0.0871, 0.4056], LM03: mean freq. difference = 0.075, 95% CI [0.0236, 0.1567]), while one genotype was more common in core compared to edge populations (LJ03: mean freq. difference = -0.191, 95% CI [-0.3334, -0.0398]). The only genotype with a significant difference between edge and core in +*Spirodela* mesocosms was LM04, which was slightly more common at the population core (mean freq. difference = -0.061, 95% CI [-0.1127,-0.0080]). While genotype-weighted trait means differed between edge and core in some replicates, these patterns were similar in mesocosms with and without competition (Fig 5B), and on average there were no differences in genotype-weighted traits between edge and core in either treatment (Table S2). Contrary to our predictions, competition did not increase leading edge genotype diversity, and there was no significant difference in inverse Simpson’s diversity index scores between treatments at the leading edge (F_1,27_ = 0.031, p=0.861), or between leading edges and cores (F_1,55_ = 1.249, p=0.269), regardless of treatment (F_1,55_ = 0.018, p = 0.894, and see Fig 5C). We found that compared to the initial populations, diversity decreased significantly in both treatments at the leading edge (mean = 1.661, 95% CI [1.418, 2.258]; CI overlapping 10 would show no significant difference from starting conditions).

**Fig 5.**
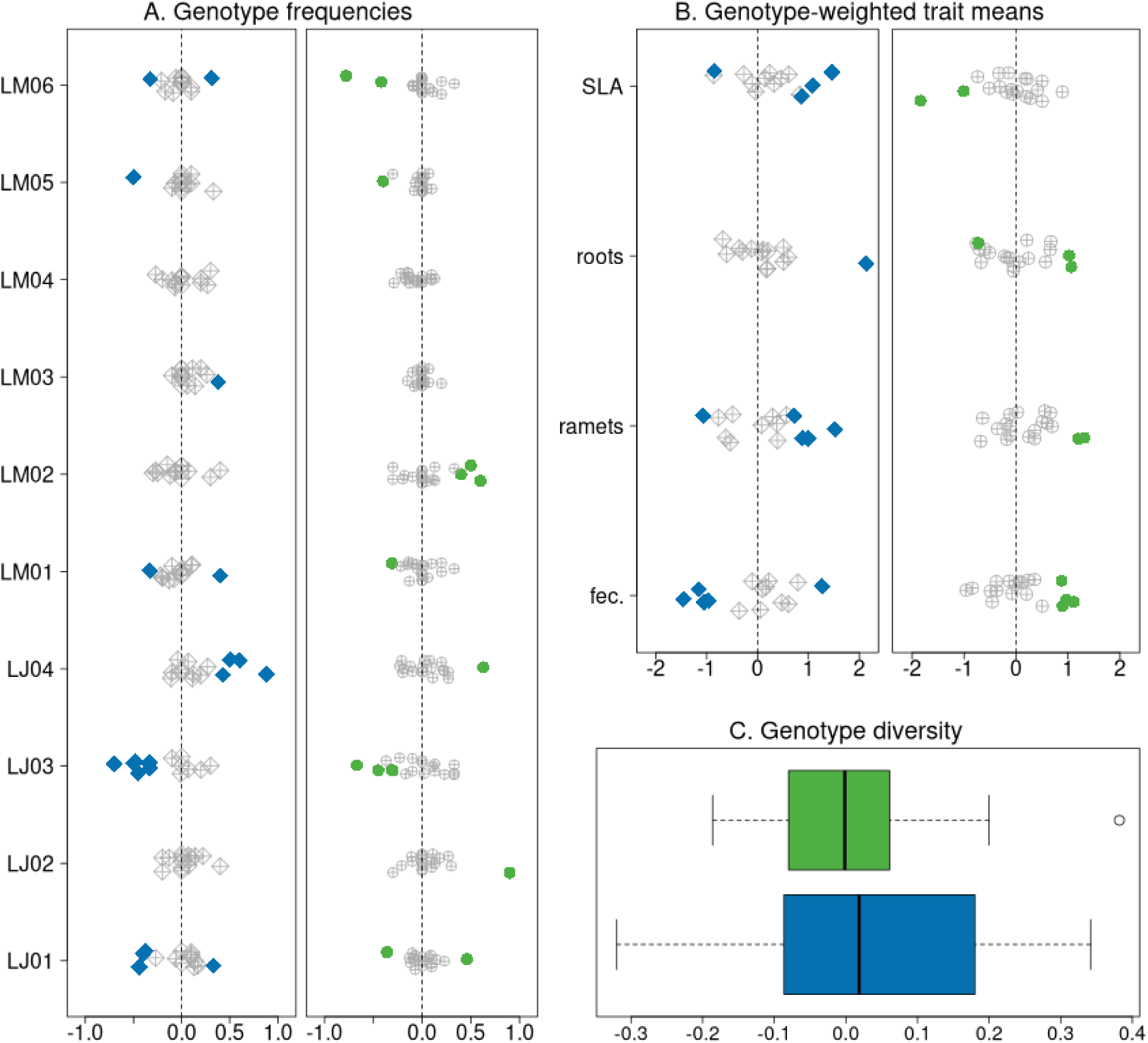
Differences in (a) genotypes (b), traits and (c) genotype diversity between the leading edge and core. (a-b) Differences in genotype frequencies (a) and genotype-weighted mean trait values (b) are shown, where one dot is the difference between edge and core for one replicate. Traits are, from top to bottom, specific leaf area (SLA), root length, (roots) number of fronts per cluster (ramet) and low density population growth rate (fec.). Replicates that were significantly different from simulated neutral distributions are shown in blue and green for *-Spirodela* and *+Spirodela* mesocosms, respectively. Grey symbols are replicates that did not have significant differences between edge and core. (c) Distributions of the loss of genotype diversity at the leading edge (compared to the core). Diversity loss is calculated as the difference in inverse Simpson’s diversity index between core and edge samples. Positive values are where genotype diversity is lower at the leading edge than at the core.

**Fig 6.**
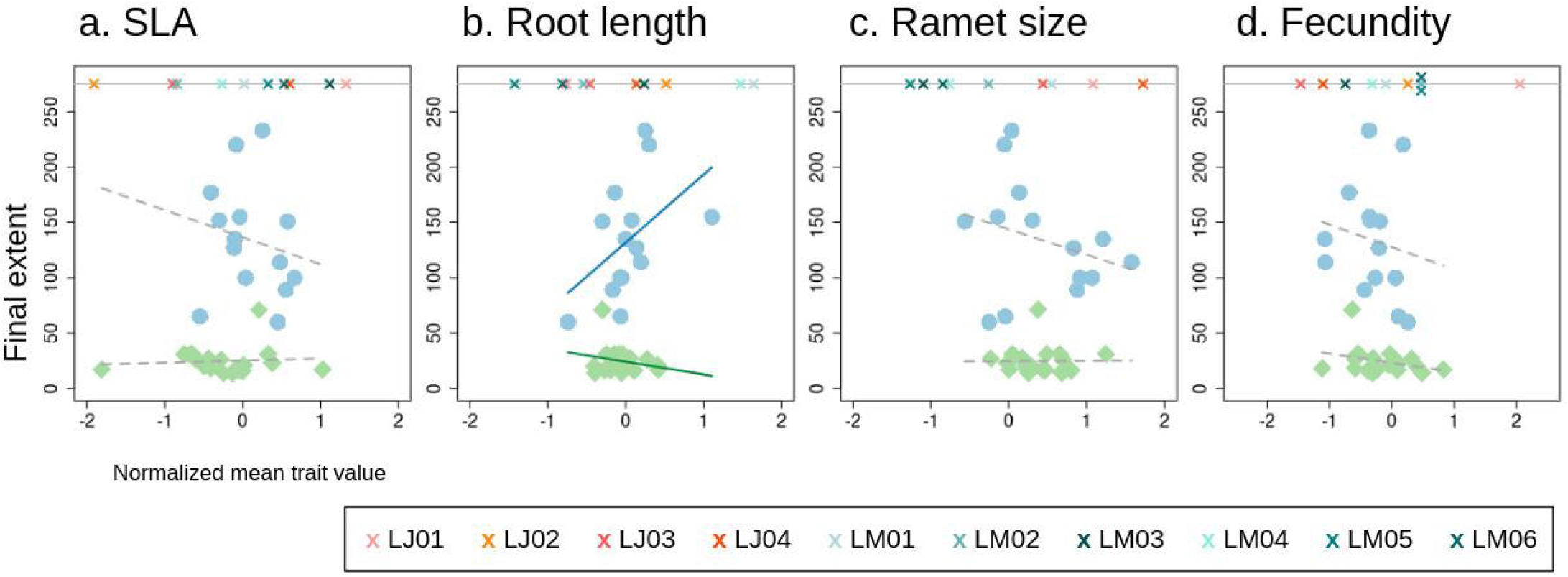
Final extent of expansions by genotype weighted trait means for each measured trait. Mesocosms with *Spirodela* are shown in green, and without in blue. Solid lines are significant modelled relationships, and dotted lines are non-significant (see Table S2). Mean normalized trait values are shown for each genotype along the top of each panel.

## Discussion

We found that competition with a resident species altered range expansion dynamics in populations of the common duckweed *Lemna*, both through evolution and demography, after fewer than 10 generations. Expanding populations in mesocosms with a resident competitor had lower absolute variability in speed, a proxy for predictability. With competition, expanding populations were significantly slower than those spreading in mesocosms without *Spirodela*. Expanding wave fronts in *+Spirodela* mesocosms were steeper, driven primarily by lower core densities as leading edge population densities did not differ between treatments. Genotype frequencies at the leading edge varied between mesocosms with and without *Spirodela*, and were more distinct from core populations in *+Spirodela* mesocosms. Contrary to our predictions, genetic diversity at the leading edge did not differ between treatments. Here we discuss our results and their role in furthering our ability to predict range expansion dynamics in the context of existing theoretical and empirical work on systems with pushed wave dynamics and evolution during range expansion.

We found that the metric we chose to measure variability (i.e., whether variability was expressed on an absolute vs. relative scale) affected the conclusion we reached about the effect of competition on variability, and thus predictability. Populations with a resident competitor had lower absolute variability in speed than those without, but higher relative variability, and only the absolute measure gave significant results. While relative metrics, such as lnCVR used here, are important to understanding ecological processes and isolating variability from differences in mean values (Nakagawa et al. 2015), we argue that they may be less appropriate when using variability as a proxy for predictability in natural populations. When making predictions about expansion speed, absolute measures can tell us directly about the range of speeds that are likely, in the appropriate unit for the process of interest. In our experiment, the higher relative variability in the +*Spirodela* treatment was largely driven by one extreme outlier in expansion extent. Without it we found lower variability with competition using both metrics. We expect that this outlier represents a rare event in nature, and we may expect that natural populations spreading into areas occupied by a resident competitor may have more predictable spread rates in general, but on rare occasions may spread much faster than expected.

Mesocosms with *Spirodela* in our experiment exhibited some features associated with pushed waves, such as steeper wave fronts, but notably did not have higher leading edge genetic diversity than was observed in *-Spirodela* mesocosms. Mathematical models and experiments demonstrating the retention of diversity with pushed waves have considered only neutral variation (Garnier and Lewis 2016; Gandhi et al. 2019; Dahirel et al. 2021), and this prediction may not hold when there are also selective processes operating at the leading edge. For example, in the same duckweed system (Usui and Angert 2024), a heat gradient reduced genotype diversity, despite the experiment’s similarities to theory that proposed retention of diversity with pushed wave dynamics (Garnier and Lewis 2016). Alternatively, it is possible that despite sharing features of pushed wave expansions, our *+Spirodela* mesocosm populations actually did not spread as pushed waves. It was not feasible in our study to test the mathematical criteria for this distinction due to a lack of knowledge about dispersal dynamics in duckweed, making it exceedingly difficult to construct meaningful dispersal kernels in order to build appropriate spread rate models. Future experiments that manipulate the wave form dynamic itself could provide better insight into the role of selection and wave form in shaping evolutionary dynamics at the leading edge.

We predicted, based on a biological definition for pushed and pulled waves (Bonnefon et al. 2013; Miller et al. 2020), that there would be less spatial genetic structure with competition than without, as more individuals would disperse from the core to the leading edge. We did find that genotype composition was more similar between edge and core in mesocosms with *Spirodela*, potentially reflecting more similar selective pressures across the expansion. Mesocosms with *Spirodela* also had shorter distances between edge and core populations, which thus increases the spatial autocorrelation between sampling locations. We observed that most populations reached equilibrium (i.e., population extents stopped increasing) within 10 days of the end of our experiment, so it is unlikely that given additional time to spread, spatial genetic structure would have emerged in mesocosms with *Spirodela*. The lack of spatial genetic structure with *Spirodela* likely reflects what we can expect to see in nature, where interspecific competition can set range boundaries and expanding populations are likely to be constrained in size and extent by resource competition (Sexton et al. 2009; Chan et al. 2019). Even if the lack of spatial genetic structure in our *+Spirodela* mesocosms is due to differences in expansion speed and extent and not the wave shape or patterns of dispersal from the edge and core, we would expect this to be generalizable beyond our experimental system.

In our experiment, we found consistent changes in some genotypes across both treatments and between core and edge, suggesting that selection was occurring. We did not find evidence for a stronger founder effect in - *Spirodela* mesocosms, instead we saw that a single genotype, LJ04, was more likely to increase in frequency than any other. Although we did not find differences in traits using genotype-weighted averages, LJ04 is the genotype with the highest mean ramet size, which may indicate increased reproductive output or dispersal ability in low stress environments (i.e. the leading edge with no competitors) and thus that some level of spatial selection was occurring without *Spirodela*. Smaller ramet sizes likely increase competitive ability in duckweeds (Zhang et al. 2020; Usui and Angert *in press*), and thus would have been more advantageous in + *Spirodela* mesocosms. While in the context of range expansions it is often assumed that neutral processes will be dominant due to short time scales and small populations, our results are less surprising when considering the broader context of evolutionary theory. That is, we should generally expect that selective processes will overcome neutral evolutionary processes where there are fitness differences between lineages.

We found that expansion in the presence of a resident species altered genotype frequencies at the leading edge, but that this did not clearly map onto the traits we measured. While we did find some evidence for selection on traits associated with competitive ability, this was observed at the leading edge and in the core in mesocosms both with and without *Spirodela.* It’s possible that selection acted on another trait we were not able to measure, or that our measured traits were not sufficient proxies for competitive ability. However, longer roots were associated with increased expansion extent in single-species mesocosms. As root length affects an individual’s ability to uptake nutrients from the water column, it likely increased fitness where nutrients were mostly widely available, i.e., empty leading edges, with increased population growth resulting in higher spread rates. As duckweed dispersal is tied closely to reproduction, we did not assess dispersal evolution, but other recent studies have shown weaker or non-existent selection for dispersal with even weak selection for other traits, such as with habitat fragmentation (Urquhart and Williams 2021; Greenbaum et al. 2022) or Allee effects (Shaw et al. 2023).

To our knowledge, only one previous experimental study conducted range expansions with a competitor, but considered two populations expanding from either end of an array (Legault et al. 2020), which may not be illustrative of the dynamics of a spreading population which encounters an established population that is at or close to equilibrium. For example, we found that steeper wave fronts with competition were largely due to a lower core population density in *+Spirodela* mesocosms, which was likely due to the presence of a resident competitor in or near the population core. Lower core population sizes were not seen in the previous experiment using flour beetles without evolution (Legault et al. 2020). There, shallower wave fronts without competition were mainly a result of higher leading edge population sizes as interspecific competition could only take place at or near the leading edge reducing population sizes here but nowhere else. The difference in experimental design – ours with a resident competitor much closer to the core of the expansion – may have reduced the carrying capacity for core populations in our experiment due to more limited resources and less physical space for the expanding species to occupy. We found that competition had a stronger limiting effect on population densities at the core than the edge, suggesting modelling competition with populations at or near equilibrium will be important to understanding competitive dynamics more likely to be seen in nature. Many predictions for pushed wave expansions are based on the expectation of higher leading edge population sizes, which would support greater genetic diversity and facilitate selection. However this pattern has not been shown in any experimental expansions with competition. Here, we did not find any difference in leading edge population size between treatments, demonstrating that higher wave steepness does not necessarily reduce founder effects or increase genetic diversity at the leading edge.

Competition in our experiment occurred for resources, largely nutrients taken up from the water column, but also for physical space. Thus, it is not surprising that many of our results align with findings from previous studies of range expansion in patchy or fragmented environments, such as slower spread and less spatial selection (Pachepsky and Levine 2011; Williams et al. 2016b, 2016a; Urquhart and Williams 2021). That higher root length increased expansion speed in mesocosms without *Spirodela*, but not those with the resident competitor, may be evidence that resource competition was an important factor in our experimental system. Longer roots may have been more advantageous beyond the leading edge of the expansion where nutrients were more abundant in empty gutters. While it’s also possible longer roots led to more dispersal by acting as rudders in the water column, we observed the opposite effect where roots become entangled and tend to anchor ramets together, instead of facilitating dispersal.

These results demonstrate the importance of conducting empirical studies that bridge theoretical predictions and conditions closer to those found in nature. Our experiment is, to our knowledge, the first to incorporate eco-evolutionary dynamics into a two-species system studying range expansion, and our design with a resident competitor more closely models natural range expansions than previous experiments assessing population dynamics. We show that biological features of pushed waves are not always correlated with each other and importantly, this research contributes to emerging evidence that even weak selective pressures may disrupt the spatial evolutionary processes often thought to be ubiquitous at the leading edge of range expansions. Experimental range expansions such as this one allow us to more accurately map eco-evolutionary theory to natural processes, and improve our ability to predict expansion dynamics in a rapidly changing world.

