## Supplemental Info for "Eco-evolutionary dynamics are shaped by competition in experimental range expansions"

### S1. DNA extractions and genotyping with microsatellite analysis

We followed the same DNA extraction and microsatellite analysis protocol as a previous paper using the same experimental system and genotypes (Usui and Angert 2025), which we describe in brief below. Duckweed tissue samples were collected, flash frozen and lysed before DNA was extracted using a modified CTAB protocol (Healey et al. 2014). We genotyped tissue samples using microsatellite analysis with four fluorescently tagged markers (Hart et al. 2019) shown on Table S1 (a). All four markers were amplified in a single 25 µL multiplex PCR reaction with one unit of 5X Taq Master Mix (New England BioLabs, Ipswich, MA), 1 µL of DNA template, and between 0.14 µM and 0.26 µM of each primer pair. The PCR protocol is described in Table S1(b). Fragment length analyses were conducted by the UBC Sequencing and Bioinformatics Consortium on an ABI 3730XL genetic analyzer (Thermo Fisher Scientific, Waltham, MA). We fit ladders and loci on each sample using Geneious Prime (2024.0.5) which were mapped to keys for each genotype developed by Usui and Angert (2024), who also delineated species using genetic barcoding.

**Table S1: Summary of microsatellite markers and PCR conditions used in genotyping**

a) Microsatellite markers and primers (developed by Hart et al. 2019) with range of fragment sizes for genotypes in this experiment

| Locus | Primer sequence F (5’–>3’) | Primer sequence R (5’–>3’) | Allele size range (bp) |
|---|---|---|---|
| R5C | TGATGCCAGTAGATCCGGC | ACGCCTGAACACGATTGATG | 320 – 380 |
| R15A | GTGACAGCGTATCCTTGTGC | TCAGCGGCAAGATCATCAAG | 220 – 280 |
| R15B | TCGAGCTAATCAGTGGAGCC | TGAGTGCTCGGCTTGACTTTC | 140 – 190 |
| R15C | TGTTCCCACCCACTTGAC | AAAGGAAGAGGGAGCAAGGG | 370 – 390 |

b) PCR conditions for single multiplex reaction

| **Thermocycler Conditions** | | | **T (°C)** | **t (sec)** |
|---|---|---|---|---|
| Initial denaturation | | | 95 | 120 |
| Denaturation | | | 94 | 30 |
| Annealing | | TD | 55 | 60 |
| Extension | **35 X** | | 68 | 30 |
| Final extension | | | 72 | 600 |

TD: touchdown, start with Annealing temp 5 ℃ higher than shown, reduce by 1 ℃ every cycle for 5 cycles until target temperature is reached.

### S2. Genotype-level trait values

We quantified genotypic variation in two traits: root length and low density fecundity, to identify selection for competitive ability? (root length) or fecundity in our experiment. Root length is expected to impact nutrient uptake (Cedergreen and Madsen, 2002), and thus we expected longer roots to be selected for with both inter- and intra-specific competition at the cores of both treatments and the leading edges with *Spirodela*. Low density fecundity has consistently been shown to be selected for at the leading edge of range expansions (Miller et al. 2020), and to strongly influence expansion speed. To measure root length, we selected 10 fronds at random for each experimental genotype from laboratory stock populations grown in the same media solution as was used in the experiment. Each root was pressed against a dark paper. Fronds were dragged upward so as to straighten the root as much as possible before measuring. Low density fecundity was measured as the per capita growth rate of single fronds grown in cups with a 50% media concentration (the same as in experimental gutters) for one week. Fecundity was measured as the number of new fronds produced in this time (roughly 1-2 generations). While we initially grew 10 replicates per genotype, one round of data collection was discarded due an error in media preparation, resulting in only 5 replicates for 3 experimental genotypes.

We supplemented this with data describing two additional traits, specific leaf area (SLA) and the number of fronds per cluster (ramet size) that are expected to be related to competitive ability  (Usui and Angert 2024). In the original study, trait data were collected across a gradient in temperatures so we retained only the replicates grown at approximately room temperature (20°C), which most closely matched ambient conditions in our greenhouse experiment.

**Table S2. Mean and confidence intervals for differences between edge and core samples within replicates**

a) Genotype frequencies

| | -*Spirodela* treatment | | | +*Spirodela* treatment | | |
|---|---|---|---|---|---|---|
| Genotype ID | Mean | 95% CI lower bound | 95% CI upper bound | Mean | 95% CI lower bound | 95% CI upper bound |
| LJ01 | -0.034 | -0.1721 | 0.0740 | 0.011 | -0.0567 | 0.0898 |
| LJ02 | 0.033 | -0.0403 | 0.1221 | 0.072 | -0.0138 | 0.2273 |
| LJ03 | -0.191 | -0.3334 | -0.0398 | -0.057 | -0.1921 | 0.0611 |
| LJ04 | 0.209 | 0.0871 | 0.4056 | 0.057 | -0.0300 | 0.1624 |
| LM01 | -0.047 | -0.1227 | 0.0642 | -0.013 | -0.0789 | 0.0560 |
| LM02 | -0.032 | -0.1171 | 0.0907 | 0.061 | -0.0342 | 0.1858 |
| LM03 | 0.075 | 0.0236 | 0.1567 | 0.004 | -0.0331 | 0.331 |
| LM04 | 0.019 | -0.0643 | 0.1078 | -0.061 | -0.1127 | -0.0080 |
| LM05 | -0.057 | -0.1198 | 0.0614 | -0.035 | -0.1099 | 0.0051 |
| LM06 | -0.025 | -0.1050 | 0.0529 | -0.033 | -0.1784 | 0.0447 |

b) Genotype-weighted mean trait values (normalized around the overall mean)

| | - *Spirodela* treatment | | | + *Spirodela* treatment | | |
|---|---|---|---|---|---|---|
| Trait | Mean | 95% CI lower bound | 95% CI upper bound | Mean | 95% CI lower bound | 95% CI upper bound |
| Root length | 0.175 | -0.0910 | 0.6606 | -0.006 | -0.2483 | 0.2658 |
| Specific leaf area | 0.278 | -0.0834 | 0.5983 | -0.107 | -0.4483 | 0.1183 |
| Ramet size | 0.168 | -0.2208 | 0.5437 | 0.215 | -0.0130 | 0.4716 |
| Low density growth rate | -0.101 | -0.5055 | 0.3128 | 0.082 | -0.1874 | 0.0824 |

**Table S3: Summary of modelled effects of traits on expansion extent**
Expansion extent as a function of genotype-weighted traits and Treatment (+ or -*Spirodela*). Each row shows the model output for the given trait plus treatment effect and interaction.

| Trait name | Trait effect | | | Treatment effect | | | Interaction | | |
|---|---|---|---|---|---|---|---|---|---|
| | coef. | t | p | coef. | t | p | coef. | t | p |
| Root length | 61.33 | 22.081 | 0.0094 | -107.736 | -9.402 | 2.63e-10 | -73.003 | -1.894 | 0.0683 |
| Specific leaf area | -24.456 | -0.975 | 0.337 | -111.416 | -8.379 | 3.10e-9 | 26.348 | 0.910 | 0.370 |
| Ramet size | -23.01 | -1.540 | 0.134 | -119.22 | -7.185 | 1.85e-13 | 23.34 | 0.842 | 0.406 |
| Low density growth | -20.24 | -0.848 | 0.404 | -104.65 | -6.969 | 1.16e-7 | 11.82 | 0.4 | 0.692 |

**S4. Sensitivity analyses for outlier in extent**
Our dataset contained one replicate in the +*Spirodela* treatment that spread considerably farther than all others, and we found that if this replicate was excluded from analyses with expansion extent, results were strongly affected. On further investigation, we did not find good reason to exclude this replicate and so chose to keep it in the dataset, while reporting results with it in the main text for analyses involving extent. Figure S1 replicates Figure 2 in the main text, with the outlier removed. Figure S2 shows the position of the outlier in the distributions of replicates for other measured attributes.This population's much further spread was driven largely by long distance spread between days 13 and 16 (24 cm spread) and 20 and 23 (21 cm spread). No maintenance actions were performed on this gutter mesocosm or any neighbouring ones between either of these dates. The only maintenance

performed on this replicate was thinning *Spirodela* density, which occurred for all +*Spirodela* mesocosms on day 10.

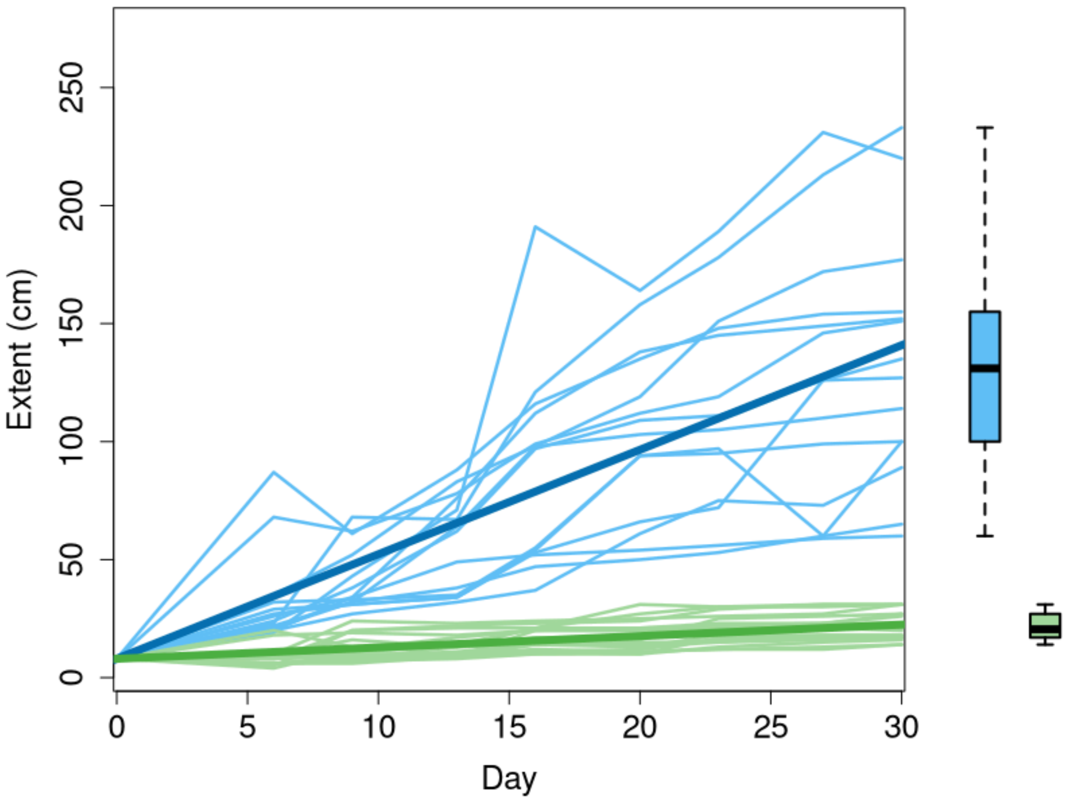


Fig S1. Copy of Figure 2 from the main text with the outlier excluded. Lines are individual replicates, darker lines are modelled extent over time for each treatment. Boxplots show the distribution of extent on Day 30. Mesocosms with and without *Spirodela* are shown in green and blue, respectively.

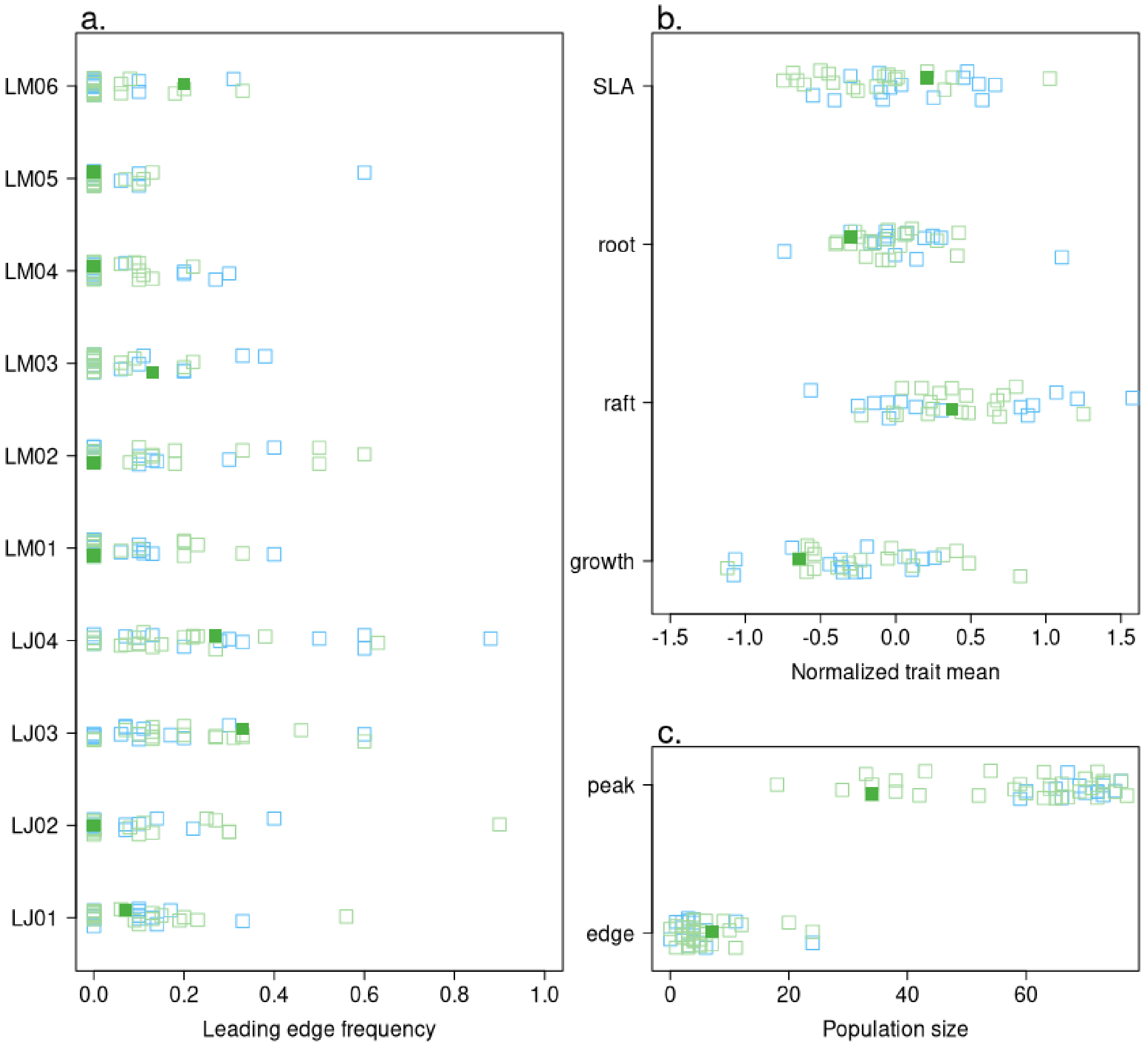


Fig S2. Outlier in extent is shown in dark green against the distributions of a. genotype frequencies and b. genotype-weighted trait means at the leading edge as well as. c. population sizes measured along transects at the edge and population core. Open squares are coloured according to competition treatment, with +*Spirodela* in green and -*Spirodela* in blue.
